# Immune Cells and Inflammatory mediators cause endothelial dysfunction in a vascular microphysiological system

**DOI:** 10.1101/2023.09.27.559626

**Authors:** Aishwarya Rengarajan, Hannah E Goldblatt, David J. Beebe, María Virumbrales-Muñoz, Derek S Boeldt

## Abstract

Functional assessment of endothelium serves as an important indicator of vascular health and is compromised in vascular disorders including hypertension, atherosclerosis, and preeclampsia. Endothelial dysfunction in these cases is linked to dysregulation of the immune system involving both changes to immune cells and increased secretion of inflammatory cytokines. Herein, we utilize a well-established microfluidic device to generate a 3-dimensional vascular Microphysiological System (MPS) consisting of a tubular blood vessel lined with Human Umbilical Vein Endothelial Cells (HUVECs) to evaluate endothelial function measured via endothelial permeability and Ca^2+^ signaling. We evaluated the effect of a mixture of factors associated with inflammation and cardiovascular disease (TNFα, VEGF-A, IL-6 at 10ng/ml each) on vascular MPS and inferred that inflammatory mediators contribute to endothelial dysfunction by disrupting the endothelial barrier over a 48-hour treatment and by diminishing coordinated Ca^2+^ activity over a 1-hour treatment.

We also evaluated the effect of peripheral blood mononuclear cells (PBMCs) on endothelial permeability and Ca^2+^ signaling in the HUVEC MPS. HUVECs were co-cultured with PBMCs either directly wherein PBMCs passed through the lumen or embedded in the supporting collagen hydrogel. We revealed that Phytohemagglutinin (PHA)-M activated PBMCs cause endothelial dysfunction in MPS both through increased permeability and decreased coordinated Ca^2+^ activity compared to non-activated PBMCs. Our MPS has potential applications in modeling cardiovascular disorders and screening for potential treatments using measures of endothelial function.

## INTRODUCTION

Endothelial cells are important contributors to the maintenance of vascular health via control of vascular permeability, maintenance of vasoactive tone, and secretion of growth factors and cytokines. Endothelial dysfunction is a common feature in cardiovascular disorders including hypertension, atherosclerosis, and preeclampsia ^1–3^. Notably, endothelial barrier function is compromised in cardiovascular diseases ^4^, atherosclerosis ^5^, and preeclampsia ^6^. Additionally, endothelial Ca^2+^ signaling is truncated in cardiovascular disorders ^7^ including hypertension ^8,9^, preeclampsia ^10^ and suggested in atherosclerosis ^11^. Ca^2+^ signaling is an important indicator of endothelial function given that Ca^2+^ is a key regulator of vasodilator production ^12^.

Cardiovascular diseases are also characterized by the presence of a proinflammatory environment that can alter endothelial function. Tumor necrosis factor (TNF) α and interleukin (IL) -6, two proinflammatory cytokines are independently correlated to endothelial dysfunction in hypertension ^13^. Additionally, vascular endothelial growth factor (VEGF) -A, classically considered an angiogenic factor, is elevated in cardiovascular diseases ^14^ and positively correlates with other inflammatory markers including IL-6 and IL-8 ^15^. TNFα, IL-6, and VEGF-A have individually shown to dysregulate Ca^2+^ signaling and barrier integrity in HUVECs (Human Umbilical Vein Endothelial Cells) in 2D cultures ^16,17^. The complex cytokine -growth factor signaling milieu in cardiovascular diseases is often underappreciated in single-factor challenge studies. Therefore, here we use an inflammatory mixture (TNFα, IL-6 and VEGF-A) on endothelial cells to overcome this.

As a major source of inflammatory cytokines, aberrant immune system involvement is implicated in multiple vascular disorders. Increased endothelial-immune interaction and changes in immune cell trafficking can increase local levels of inflammatory cytokines at the endothelium ^18–20^. Indeed, activation of immune cells and increased interaction of immune cells with vasculature is known to occur in hypertension ^21^. Further, previous studies have shown that immune cells are implicated in negatively regulating endothelial function by causing decreased Ca^2+^ signaling ^22^, changing adhesion molecule expression, and allowing for immune cell transmigration ^23^. We therefore utilize peripheral blood mononuclear cells (PBMCs) with or without activation and study their effects on endothelial cells in this study.

Traditionally, *in vitro* analyses using HUVECs rely on 2-dimensional (2D) cultures in flat polystyrene plates or transwells. However, a drawback of traditional 2D *in vitro* models is that the in vivo tubular organization of endothelial cells, which is key to their function and secretory profile ^24^, cannot be recapitulated. Further, key autocrine and paracrine signaling in these interactions is diluted in the larger volumes of 2D polystyrene plates. To overcome these issues, there is recent increased interest in recapitulating structure of tubular organs generally ^25^ and specifically with respect to studying vascular pathophysiology ^26^ and cardiovascular disorders ^27^. We therefore chose to leverage an existing tubular endothelial vessel microphysiological system (MPS) to model endothelial dysfunction ^28^. Human Umbilical Vein Endothelial Cells (HUVECs) are easy to isolate and a well-established cell model to study cardiovascular diseases ^29^, making them a readily available candidate for evaluating and characterizing endothelial function in our MPS of choice. Our MPS system also offers the flexibility to be used with immune cell co-cultures, key to deciphering the effect of immune cell crosstalk on endothelial cell function.

Here, we assessed the impact of inflammatory mediators on endothelial function using either an inflammatory mixture (i.e., TNFα, IL-6 and VEGF-A) or PBMCs on vascular MPS, in order to quantify endothelial function in response to an inflammatory microenvironment. We assessed endothelial function by measuring endothelial barrier function and Ca^2+^ signaling sequentially and characterizing endothelial-immune crosstalk via assessment of cytokine secretions in the experimental media. We show herein that both the inflammatory mixture and co-culture with activated PBMCs alter endothelial monolayer integrity and Ca^2+^ co-activity, thus demonstrating broad endothelial dysfunction in our HUVEC MPS. We also observe increased inflammatory cytokine secretions in our MPS concurrent with endothelial dysfunction. Importantly, our MPS recapitulates in vivo Ca^2+^ signaling of intact vessels *in vitro*, thus more closely recapitulating vascular function than traditional 2D models, and laying the ground for translational studies on cardiovascular diseases.

## METHODS

### Human cell/tissue collection

Human Umbilical Vein Endothelial Cells were obtained from umbilical cords of normotensive women, collected at Meriter Hospital, Madison, WI and jointly approved by Institutional Review Board (IRB) at University of Wisconsin, Madison and Meriter Hospital, Madison. Consent was obtained for tissue collection and samples were deidentified. Collagenase B solution 2 mg ml^−1^ B (Roche, Indiana) was used for HUVEC isolation (detailed protocol in ^10^). HUVECs isolated from 5 individuals were pooled at passage 3 and frozen. Pooled HUVECs were expanded in a T75 flask in ECM media (Endothelial cell medium, ScienCell) consisting of 500 mL basal media, 25 mL fetal bovine serum, 5 mL ECGS. Cells were grown to ∼90% confluence and routinely trypsinized with 0.05% trypsin EDTA (Gibco, Massachusetts).

Whole blood for peripheral blood mononuclear cell (PBMC) collection was obtained from American Red Cross, Madison. Blood collection for PBMC isolation was given a waiver from the IRB as ‘Not Humans Subject Research’. PBMCs were isolated using gradient centrifugation with Ficoll paque plus (Cytiva, Massachusetts).

### Microdevice fabrication

Microdevices were fabricated as previously described ^28^ in PDMS (polydimethylsiloxane) using conventional soft lithography. This device consists of two PDMS layers that define a chamber and sandwich a PDMS rod that is suspended between them. Briefly, photomasks were designed using Illustrator software, and used to generate SU-8 master molds. PDMS (10:1 base polymer to curing agent ratio) was poured on top of the SU-8 master molds and baked at 80 ºC for 4 h. Then, PDMS layers were manually aligned. PDMS rods were fabricated from 23-gauge hypodermic needles (inner diameter -230 μm, 14-840-88, Fisher scientific, Pittsburgh, PA), extracted from the needles, and inserted between the layers using fine tweezers. This device was then oxygen-plasma bonded (Diener Femto system) onto 60 mm glass bottom dishes (Mattek Corporation).

### Collagen hydrogel preparation

Prior to cell seeding, the dishes were UV-sterilized for 15 minutes. Devices were then treated with 2% poly(ethyleneimine) (PEI, Sigma-Aldrich, 03880) and 0.4% glutaraldehyde (Sigma-Aldrich, G6257) in distilled water for 10 and 20 min respectively for improving attachment of hydrogel to the device. The devices were then washed with cell-culture grade water 4 times, and thoroughly dried. A Rat tail Collagen type I hydrogel was prepared (Corning, 354249) at a final concentration of 4.5 mg ml^−1^ and pH of 7.2. This hydrogel solution also consisted of 30 µg ml^−1^ Fibronectin (Sigma-Aldrich, F1141), 0.5 M NaOH (Fisher Scientific, S318), 10X PBS and cell culture media ECM (Sciencell, 1001) (details in table 1). Collagen hydrogel (6 μl per device) was then pipetted through the side ports and allowed to polymerize for 30 min. The PDMS rod was removed with an isopropanol-sterilized tweezer. The resulting tubular structure was washed with ECM media three times before cell seeding. In MPS containing embedded PBMCs, PBMCs were used at 4*10^6^ cells ml^−1^, roughly equating to 2000 cells in the collagen hydrogel.

**Table 1:**
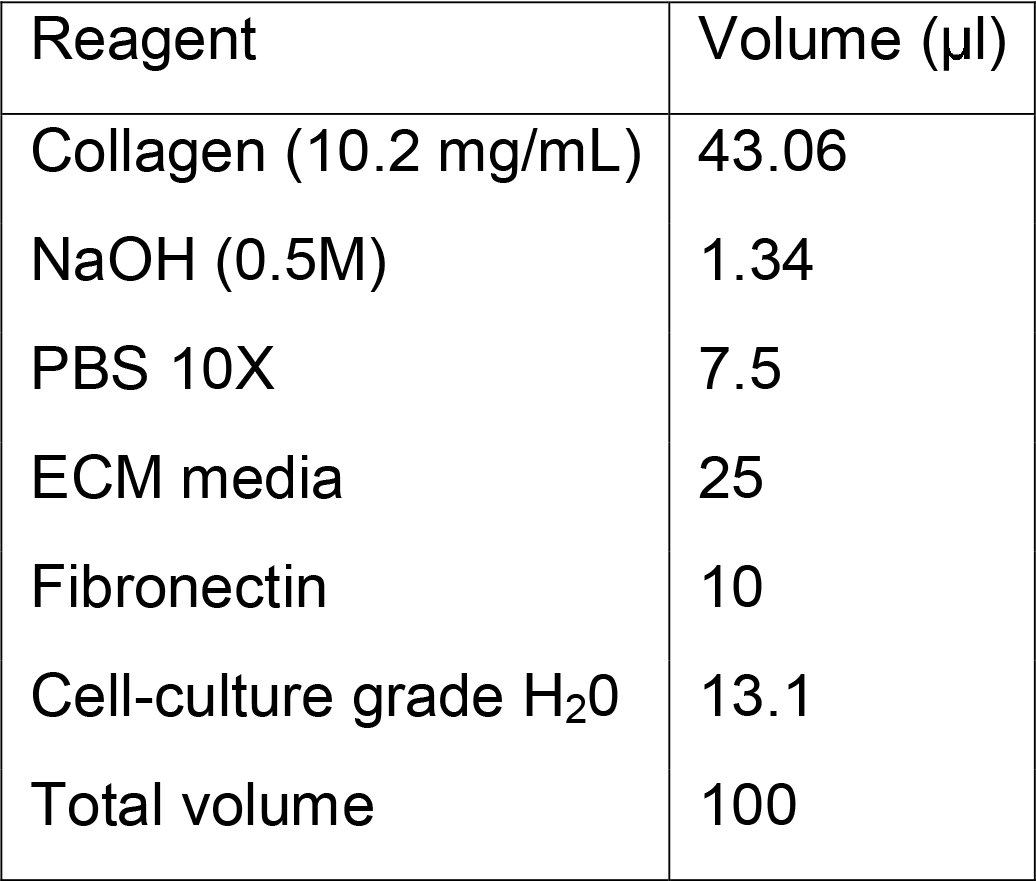
Recipe for collagen hydrogel.

### Cell culture in microdevices

Trypzinized HUVECs were resuspended at a concentration of 20,000 cells μl^−1^. Approximately 2 μl of cell suspension was added to each lumen and cells were allowed to adhere for 1 h at 37°C. Excess cells were washed with ECM media thereafter. Cells were maintained for 3 days, following which permeability and Ca^2+^ imaging assays were carried out.

MPSs were conditioned with an inflammatory mixture that consisted of TNFα, VEGF and IL-6 at 10 ng/ml each. Lumenal PBMCs were used at 2,000 cells μl^−1^ in ECM media to maintain 1:1 ratio of PBMCs:HUVECs. Lumenal Act PBMCs were activated with Phytohemagglutinin-M 10 µg ml^−1^ (Roche, Indiana), with cell concentration maintained at 2,000 cells μl^−1^.

### Immunofluorescence protocol

MPSs were fixed with 4% paraformaldehyde (PFA) (EMScience, 15700) for 15 min. MPS was incubated with blocking buffer (3% BSA in 0.1% PBS-Tween 80) over two nights. MPS was washed three times with washing buffer (0.1% PBS-Tween 80). This was followed by incubation with primary antibodies at 4○C overnight. Primary antibodies were diluted at 1:50 for ms CD31 and 1:50 for rb VE-cadherin. After rigorous washing, MPS was incubated with secondary antibodies (Goat anti-mouse IgG 488 and Goat anti-rabbit IgG 647) and nucleic dye (Hoechst 33342) (1:1000 concentration for all). Confocal images were acquired using a Leica SP8 3× STED super-resolution microscope (Wetzlar, Germany) in the UW-Madison Optical Imaging Core. Images were processed using ImageJ.

### Permeability quantification

After 3 days of culture, permeability of the vascular MPS was assessed by adding 1-2 μl of 1 μM dextran 70 kDa to the lumen. Images were taken every 5 min for 15 min on a Nikon Ti-Eclipse microscope using 6X magnification using the Texas Red filters. Diffusion of the dextran dye outside the lumen formed the basis for calculating permeability, measured via image intensity using Image J. Specifically, the following formula was utilized for calculating permeability ^30^:

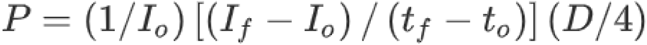

where I_0_ is the total initial intensity outside the lumen, I_f_ is the total intensity outside the lumen at 15 min, t_o_ is the initial time point, t_f_ is the final time point of 15 min, and D is lumen diameter.

### Ca^2+^ imaging

MPSs were perfused twice with Fluo-8 AM dye 10 μM and incubated for 30 min at 37°C. MPSs were then washed with KREBS buffer (125 mM NaCl, 5 mM KCl, 1 mM MgSO_4_, 1 mM KH_2_PO_4_, 6 mM glucose, 2 mM CaCl_2_, 25 mM HEPES, pH = 7.4) 4 times and allowed to hydrolyze for 30 min at room temperature. MPSs were then imaged for 10 min with 1 image sec^−1^ with emission wavelength at 495 nm on the same microscope. 100 μM ATP was added within the first few seconds of these recordings.

### Ca^2+^ network analysis

Cells in the MPS were manually segmented using Image J. The MATLAB program FluoroSNNAP ^31^ was subsequently used to measure Ca^2+^ co-activity. Specifically, the following steps were carried out in the program – quantification of fluorescence for each cell with time, quantification of change in fluorescence (ΔF) with respect to initial fluorescence for each cell with time, identification of Ca^2+^ events defined where (ΔF/F > 0.01) for each timepoint. Subsequently, time points showing the most events were identified and a collection of cells active at these time points were chosen as belonging to a core ensemble ^32^. The percentage of cells that were active (i.e. showing Ca^2+^ events at a given time point) within the ensemble were then computed in FluoroSNNAP.

### Conditioned media analysis

Conditioned media was collected from HUVEC MPSs and assayed using LXSAHM-16 (R&D Systems) and HCYTOMAG 41 cytokine (MilliporeSigma) panels. Media from at least 2 experimental replicates were used for analysis, and at least 2 technical replicates were included for each condition according to manufacturer’s instructions and assayed on Luminex 200. Luminex xMAP or Belysa software were utilized for making standard curves and identifying concentrations.

### HUVEC culturing, Ca^2+^ Imaging and analysis for 2D culture controls

HUVECs were grown in ECM media in clear wells for ∼4-5 days until confluence. The cells were loaded with Fluo-8 (5 μM) for 40 min in the incubator followed by 30 minutes of hydrolysis in KREBS buffer at room temperature. Cells were imaged on Biotek Cytation 5 using the GFP filter with excitation/emission at 488nm/510nm. Cells were segmented using Cell Profiler ^33^. The rest of the Ca^2+^ analysis methodology was the same as for 3D culture.

### Statistics and replicates

Statistics were performed using t-test for permeability, Ca^2+^ co-activity and growth factor or cytokine concentrations. n≥5 MPSs were used for each condition for permeability and Ca^2+^ co-activity analysis.

## RESULTS

### Development of a HUVEC MPS model

We report a vascular MPS model exposed to different sources of inflammation and evaluate subsequent effects on endothelial function. To develop this model, we used an existing and well-validated microfluidic device that generates 3D tubular blood vessel models (fig 1d). We used Human Umbilical Vein Endothelial Cells which express relevant markers CD31 and VE-cadherin (fig 1b, 1c). We subsequently leverage this vascular MPS model to evaluate macrovascular endothelial function as a model of cardiovascular disease and associated persistent inflammation derived from inflammatory cytokines or activated immune cells. To this end, we characterized endothelial function using metrics of barrier function (measured as vascular permeability to fluorescent dextran) and Ca^2+^ signaling in the following result sections.

**Fig 1.**
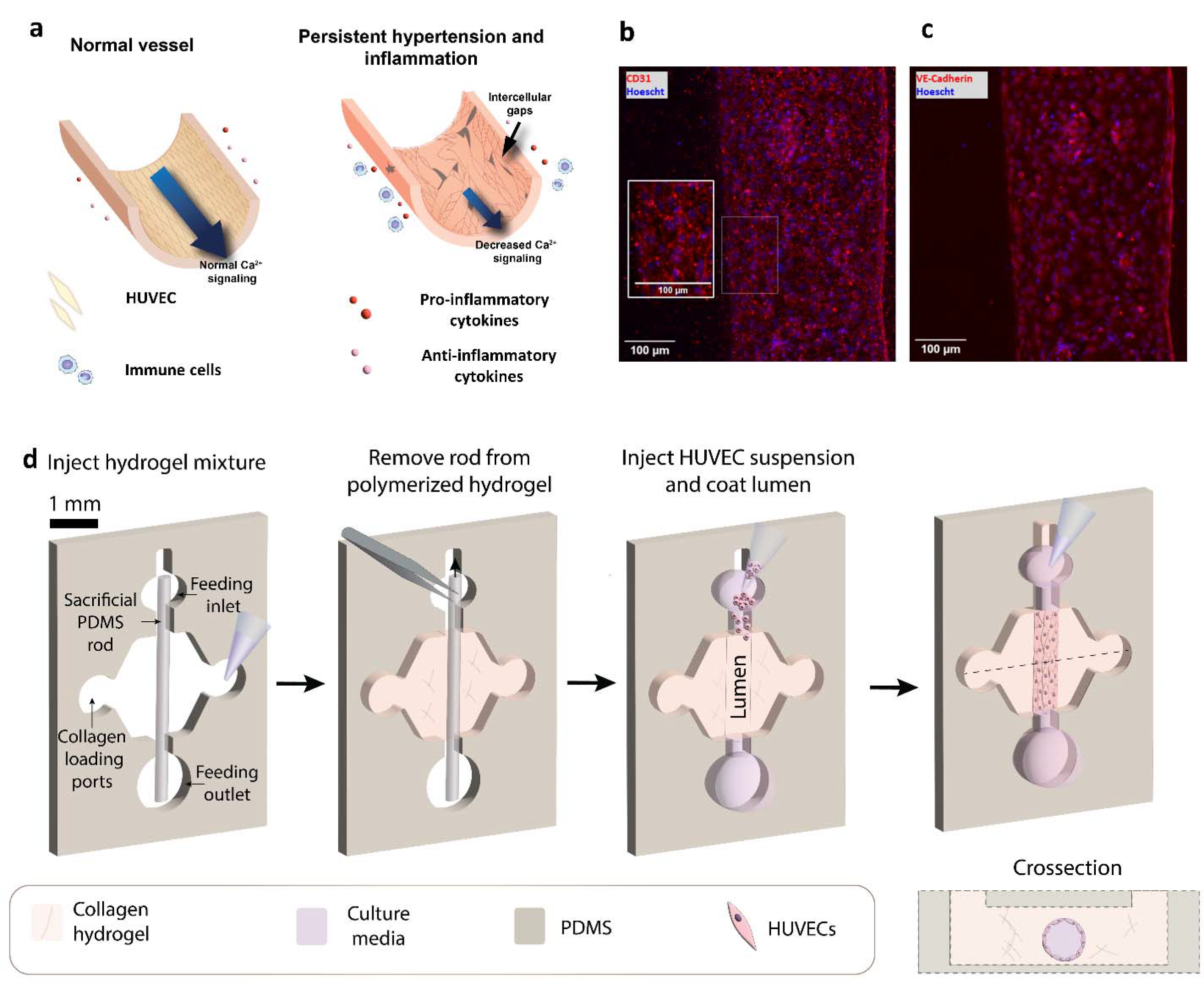
HUVEC MPS preparation and staining for endothelial markers. a - Schematic demonstrating normal vessel and vessels exposed to inflammatory cytokines or PBMCs. b,c - Staining of HUVEC MPS with endothelial marker CD31 and junctional protein VE-cadherin. Nucleus stained with Hoechst 33342. d - Schematic depicting the structure of the MPS device and the steps involved in MPS preparation.

### Inflammatory mixture causes increased permeability in vascular MPS

Endothelial monolayer permeability was assessed by diffusion tracking of 70kDa dextran dye (fig 2a). HUVEC MPS displays barrier properties (fig 2 b,c) as inferred by observed permeability of 2.81 × 10^−5^ ± 2.68 × 10^−6^ cm s^-1^, compared to cell-free MPS (5.11 × 10^−5^ ± 3.12 × 10^−6^ cm s^-1^). HUVEC MPS showed significantly higher permeability upon long-term (48 h) treatment with inflammatory mixture (TNFα, IL-6, VEGF - 10ng/ml each - termed ‘mix 48h’) at 4.79 × 10^−5^ ± 5.81 × 10^−6^ cm/s compared to control MPS; and showed no difference from cell-free MPS.

**Fig 2.**
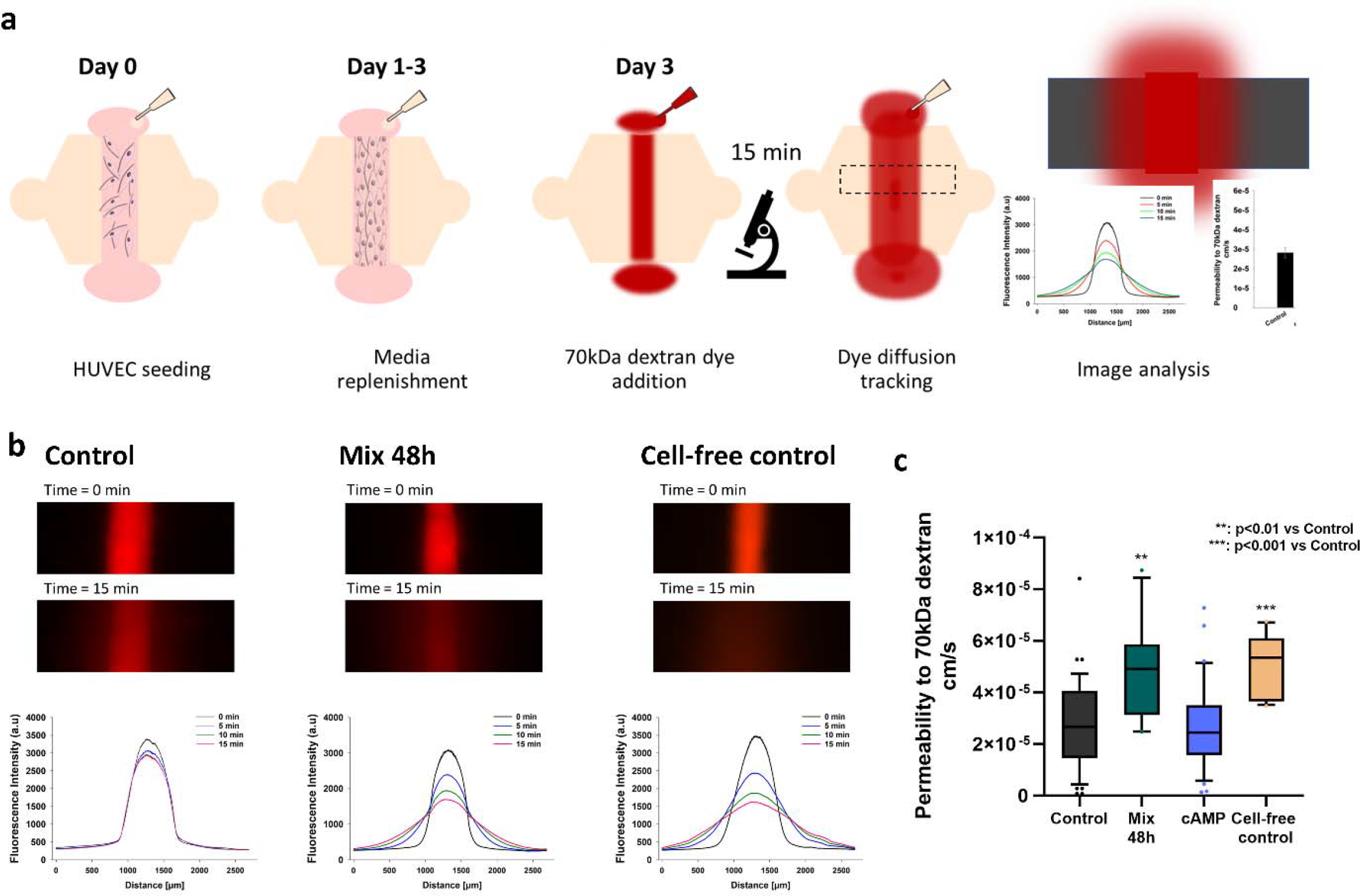
HUVEC MPS show increased permeability upon treatment with inflammatory cocktail (mix 48h). a - Schematic shows steps involved in measuring permeability of HUVEC MPS using diffusion of a 70kDa fluorescent dye. MPS was loaded with 1 µM 70kDa dextran dye and images were taken every 5 minutes for 15 minutes to track diffusion of dye. b - Fluorescence intensity plots and images of HUVEC MPS upon addition of 70kDa dextran red. Representative fluorescent images and fluorescence intensity plots are included for different conditions. c - Box plots for permeability data are plotted for HUVEC MPS under different treatments including the inflammatory cocktail. n≥5, statistics was performed using t- test.

### Inflammatory mixture stimulates secretion of angiogenic factors in HUVEC MPS

HUVEC MPS exposed to the inflammatory mixture led to increased secretion of G-CSF (Granulocyte-colony stimulating factor) and uPA (urokinase-type plasminogen activator) (fig 3b). Increase in VEGF-C was also observed in HUVEC MPS exposed to the inflammatory mixture (fig 3a). Additionally, chemokine ligands and receptors were significantly increased in MPS treated with the inflammatory mixture (156-fold increase in CCL-5, 7-fold increase in CCL-7, 32-fold increase in CX3CL1 and 120-fold increase in CXCL10) as shown in (supp tables 1,2).

**Fig 3.**
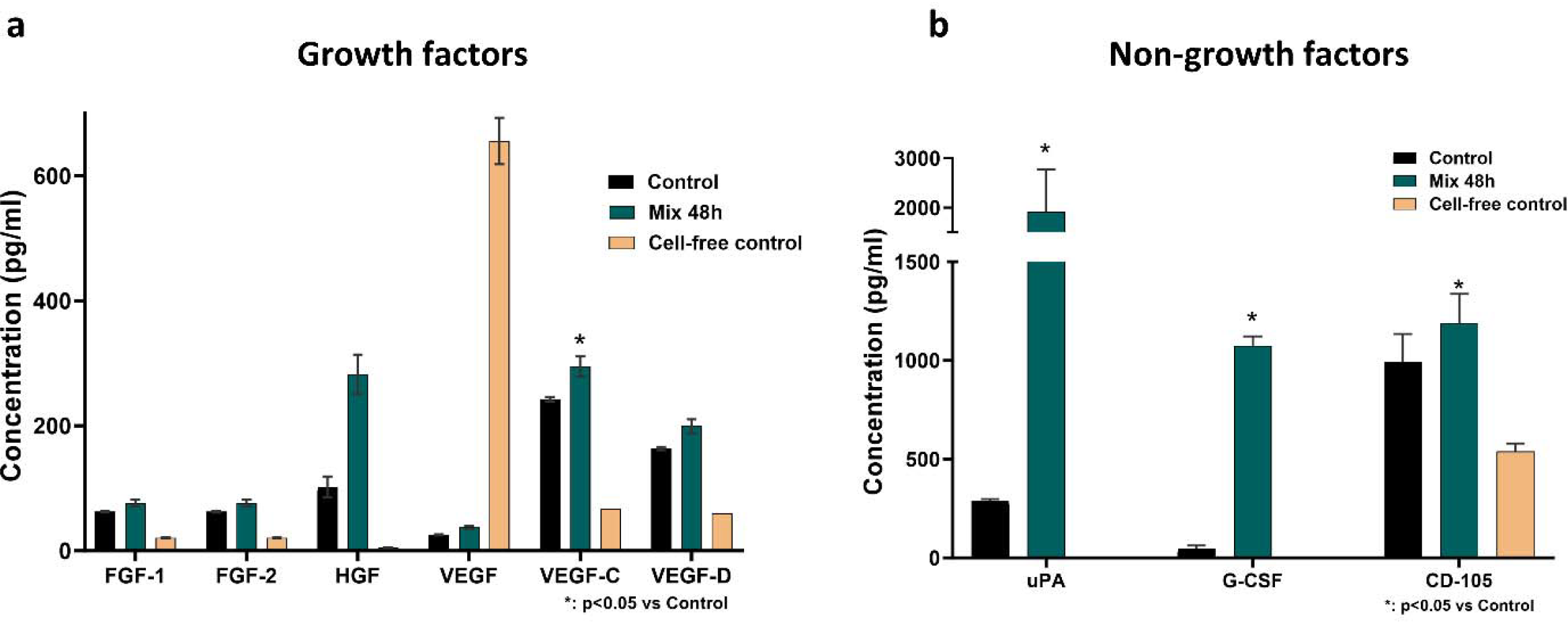
Inflammatory cocktail leads to increased secretion of VEGF-C, uPA and G- CSF in HUVEC MPS. a - Graph shows average concentration of growth factors in conditioned media collected from HUVEC MPS (control and mix). Statistics was performed using t-test. b - Graph shows average concentration of non-growth factors in conditioned media collected from HUVEC MPS. Statistics was performed using t-test.

### HUVEC MPS treated with the inflammatory mixture show reduced Ca^2+^ co-activity

We leveraged an advanced programming tool, FluoroSNNAP, to automatically measure coordinated activity within a HUVEC MPS. Co-activity refers to the percentage of cells within a core ensemble of cells that show Ca^2+^ events at a time point (fig 4a). Both HUVEC MPS and HUVECs in 2D culture display an initial co-activity peak in 90-100% of cells upon stimulation with 100 μM ATP indicating the viability of cells and a near perfect coordinated Ca^2+^ release in response to ATP. Control MPS continues to display a high level of sustained Ca^2+^ co-activity (70-75%) 4-10 mins after ATP stimulation. However, HUVECs in control 2D show low levels of Ca^2+^ co-activity (Fig 4b). Typical tracings for individual lumens are shown (fig 4c-e).

**Fig 4.**
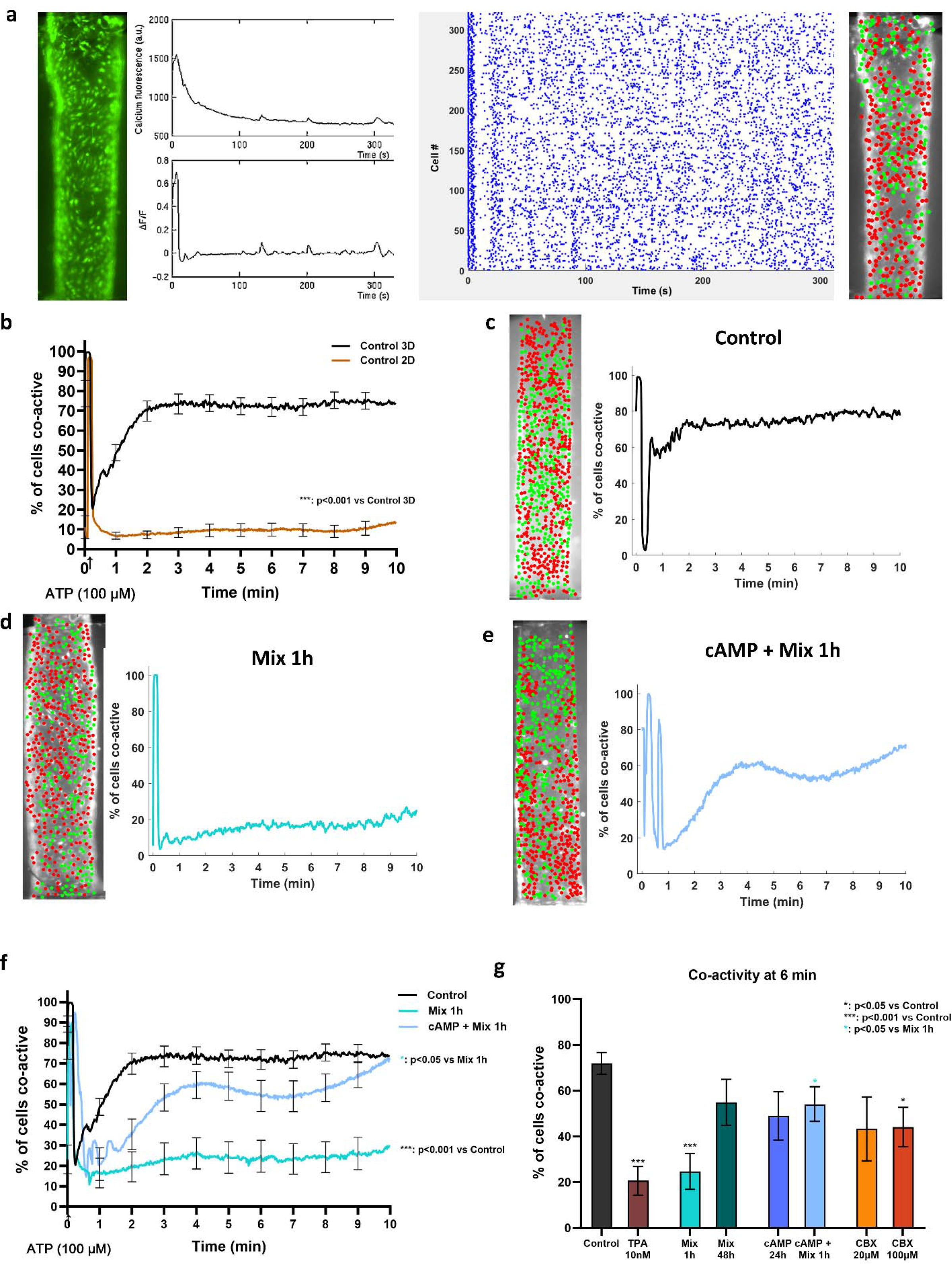
Ca^2+^ co-activity in HUVEC MPS is decreased by the inflammatory cocktail. a - Schematic demonstrating the analysis pipeline in FluoroSNNAP. Cells loaded with Fluo-8 (Ca^2+^ indicator) are shown. Fluorescence analysis on a single cell using FluoroSNNAP is shown with time, followed by ΔF/F analysis with time. FluoroSNNAP generated raster plot with all identified events (ΔF/F>0.01) shown for the first 5 minutes. Coactive frames from the raster plot are used to identify a core ensemble of cells (green) used for subsequent analysis. b - Comparison of co-activity of cells belonging to the core ensemble between HUVEC MPS (3D culture) and HUVECs on 2D conventional tissue culture plates demonstrates high co-activity in HUVEC MPS. Statistics was performed using t-test using area under the curve measurement. c,d,e - Representative images of core ensemble and co-activity plots from a single MPS in different conditions.Control, Mix 1h and cAMP + mix 1h are included to demonstrate reduction and recovery in co-activity occur with mix 1h and cAMP + mix 1h respectively. f - Average co-activity of HUVEC MPS in different conditions is shown here. Control MPS shows the highest co-activity. Statistics was performed using t-test using area under the curve measurement. g - Mean co-activity at 6 mins computed from HUVEC MPS in response to different treatments is shown here. TPA and CBX negatively affect gap junction communication and cause decreased co-activity in HUVEC MPS. n≥5, statistics was performed using t-test on co-activity value at 6 min.

Ca^2+^ signaling in HUVECs is dependent on cell-cell communication through gap junctions, primarily via Cx43 ^34^. Cells were therefore treated with TPA (12-O-Tetradecanoylphorbol-13-acetate), a gap junction inhibitor that causes inhibitory phosphorylation on Cx43 ^35^; carbenoxolone (CBX), a gap junction blocker ^36^; or cAMP a PKA agonist, involved in gap junction assembly and membrane trafficking ^37^. Mean tracings for control, inflammatory mix, and cAMP are shown (fig 4f). Quantification of co-activity for all treatments is shown in fig 4g. HUVEC MPS treated with TPA 10 nM show 20-30% co-activity at 4-10 mins, decreased significantly from control MPS. MPS treated with inflammatory mixture for 1h also show a similar 20-30% co-activity profile, also decreased from control MPS. However, MPS treated with the inflammatory mixture for 48h show 55-63% co-activity and failed to show any difference in comparison to control MPS, while showing significant improvement in comparison to the inflammatory mixture for 1 h. Treatment with cAMP 1 mM (24 h) did not change co-activity compared to control MPS. However, cAMP 24 h treatment followed by addition of the inflammatory mixture for 1 h offered significant improvement in Ca^2+^ co-activity when compared to MPS treated with the inflammatory mixture for 1h only. CBX did not cause any change in co-activity at 20 μM, while CBX 100 μM showed a significant reduction in co-activity compared to control MPS.

### PBMCs in HUVEC MPS alters endothelial monolayer permeability

Two methods of PBMC introduction were utilized to mimic signaling from immune cells embedded in the vascular wall (embedded PBMCs) or from the vascular lumen (lumenal PBMCs) (fig 5a). HUVEC MPS with embedded PBMCs (PBMCs embedded in collagen hydrogel surrounding the MPS) revealed increased barrier function compared to control MPS. Further, MPS treated with lumenal PBMCs (PBMCs passing through the MPS) for 24h showed no change in permeability against control MPS. Interestingly, MPS treated with lumenal activated PBMCs for 24 h (10ug/ml PHA-M activation) showed significantly increased permeability in comparison to MPS treated with lumenal PBMCs (fig 5b).

**Fig 5.**
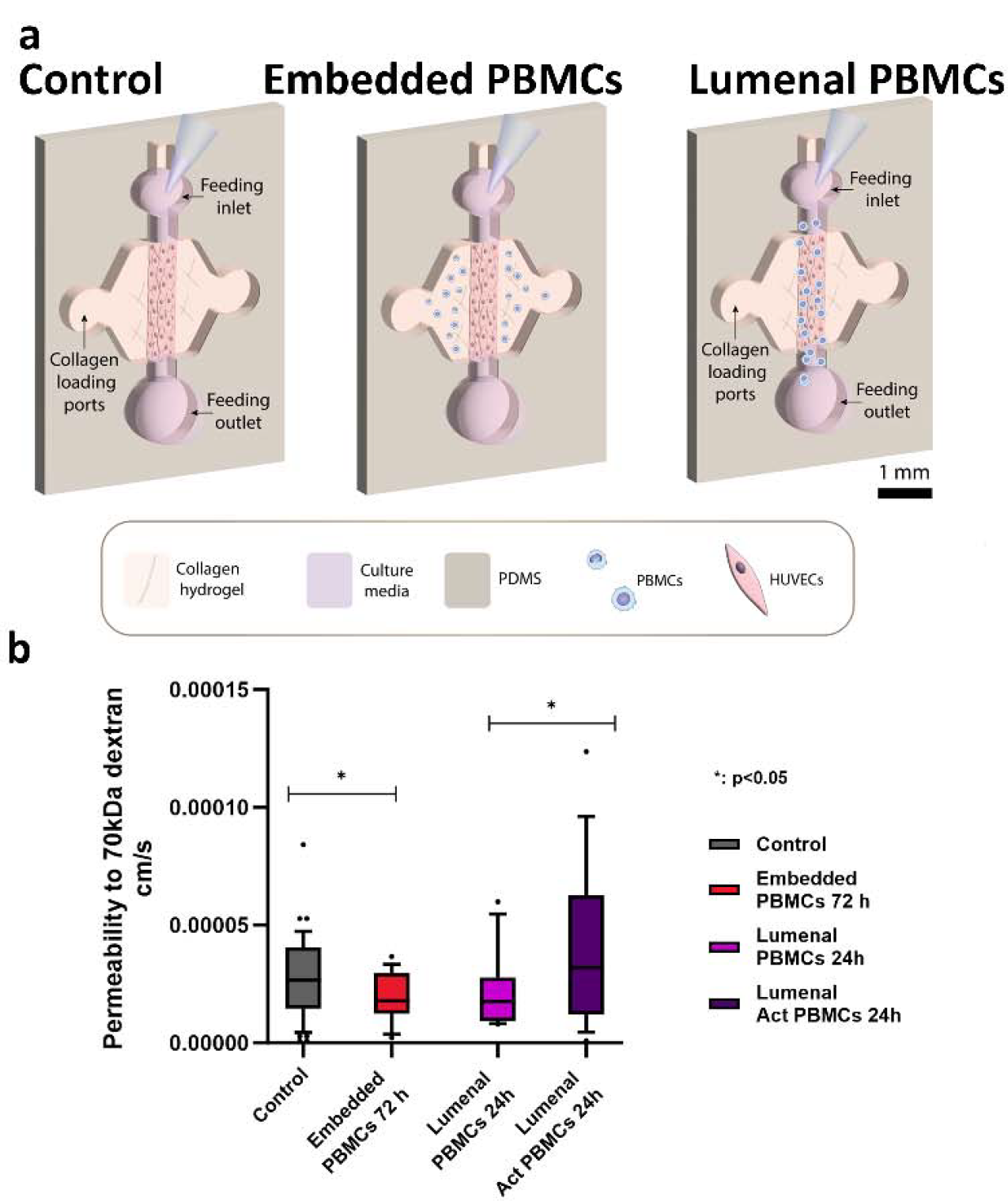
Activated PBMCs directly interacting with HUVEC MPS show increased permeability compared to non-activated PBMCs interacting with HUVEC MPS. a - Schematic depicting the HUVEC MPS interacting with PBMCs. Embedded PBMCs indirectly interact with HUVEC MPS while lumenal PBMCs directly interact with the HUVEC MPS. b - Box plots for permeability data are plotted for HUVEC MPS under different types of PBMC interactions. n≥5, statistics was performed using t-test.

### Co-culture of HUVEC MPS with PBMCs results in decreased Ca^2+^ co-activity

HUVEC MPS with embedded PBMCs sorted into two categories of responses; either they showed low-coactivity similar to the profiles from inflammatory mixture 1h MPS or they show high co-activity similar to the profiles from control MPS (fig 6a). Overall, Embedded PBMCs showed a mean of 38-45% co-activity, significantly lower than control MPS (fig 6c,d). In MPS with lumenal PBMCs a single co-activity profile was apparent (fig 6b). Overall, with lumenal PBMCs, Ca^2+^co-activity showed a trend towards decrease (p=0.054 based on area under co-activity curve) in comparison to control MPS (fig 6 c,d). MPS with lumenal activated PBMCs showed significantly lower Ca^2+^ co-activity in contrast with PBMCs through the MPS or control MPS (fig 6c,d).

**Fig 6.**
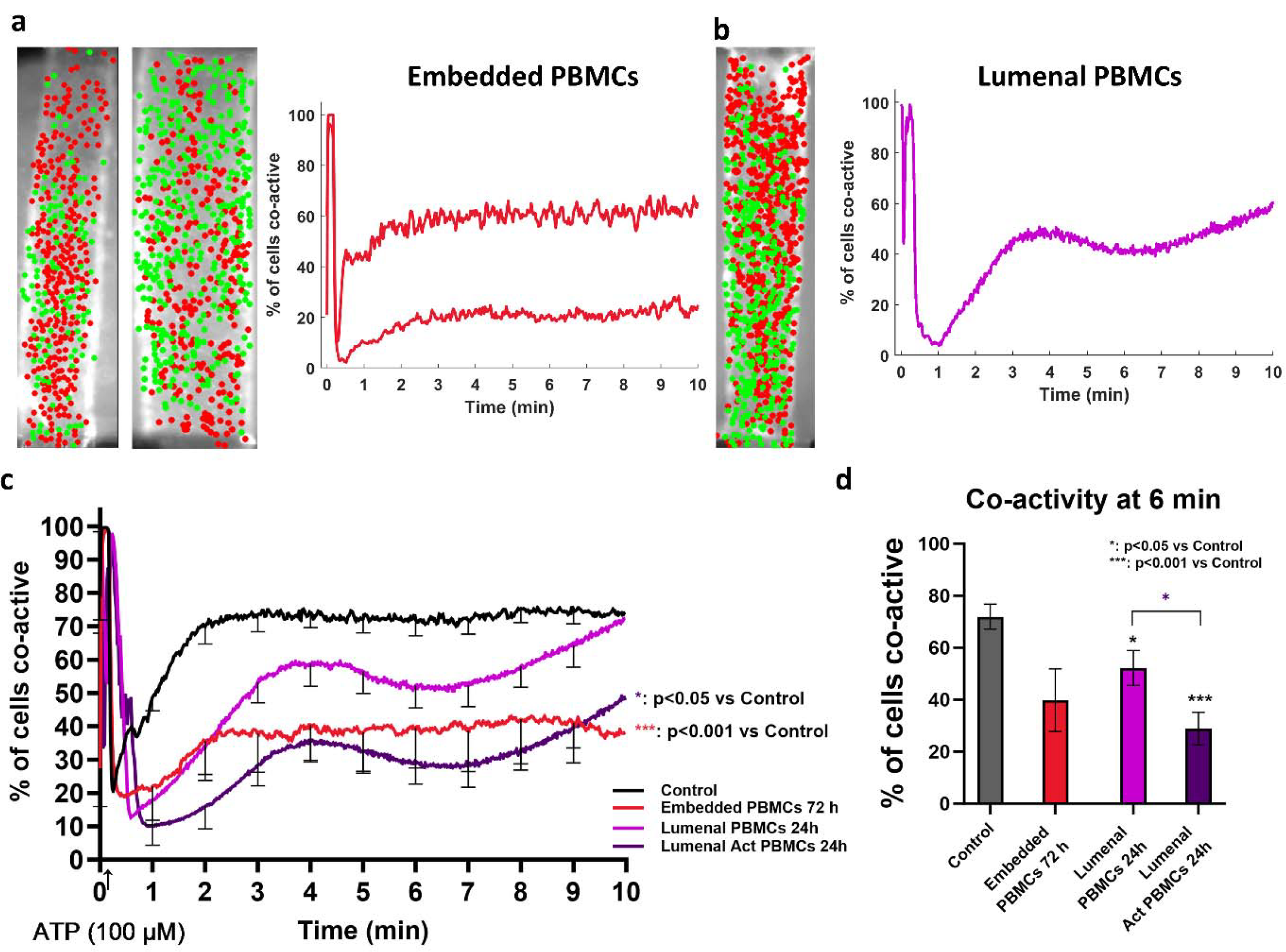
Coordinated Ca^2+^ activity is decreased in HUVEC MPS exposed to lumenal PBMCs with or without activation. a - Representative image of core ensemble and co-activity plot from two different MPSs with embedded PBMCs. b - Representative image of core ensemble and co-activity plot from one MPS with lumenal PBMCs. c - Average co-activity of HUVEC MPS interacting with PBMCs is shown here. Control MPS shows the highest co-activity. n≥5, statistics was performed using t-test using area under the curve measurement. d - Mean co-activity at 6 mins computed from HUVEC MPS interacting with PBMCs. n≥5, statistics was performed using t-test on co-activity value at 6 min.

### Activated PBMCs lead to increased secretion of cytokines and chemokines in HUVEC MPS - PBMC co-culture

In order to assess the mixture of inflammatory mediators secreted in the MPS and therefore available to the HUVECs, we assayed the experimental media at the termination of the experiment and subjected it to multiplex assay. Analysis of media from MPS treated with lumenal activated PBMCs revealed increased secretion of inflammatory cytokines, including interleukins (IL-1a, IL-1b, IL-2. IL-10), IFNγ and TNFα (fig 7a). Growth factors and colony stimulating factors were largely unchanged (Fig 7b). Additionally, chemokines including CCL-3, CCL-5, CCL-7 and IP-10 (CXCL10) showed ≥2 fold increase in lumenal activated PBMC-MPS co-culture compared to lumenal PBMC-MPS co-culture (fig 7c, supp tables 1,2).

**Fig 7.**
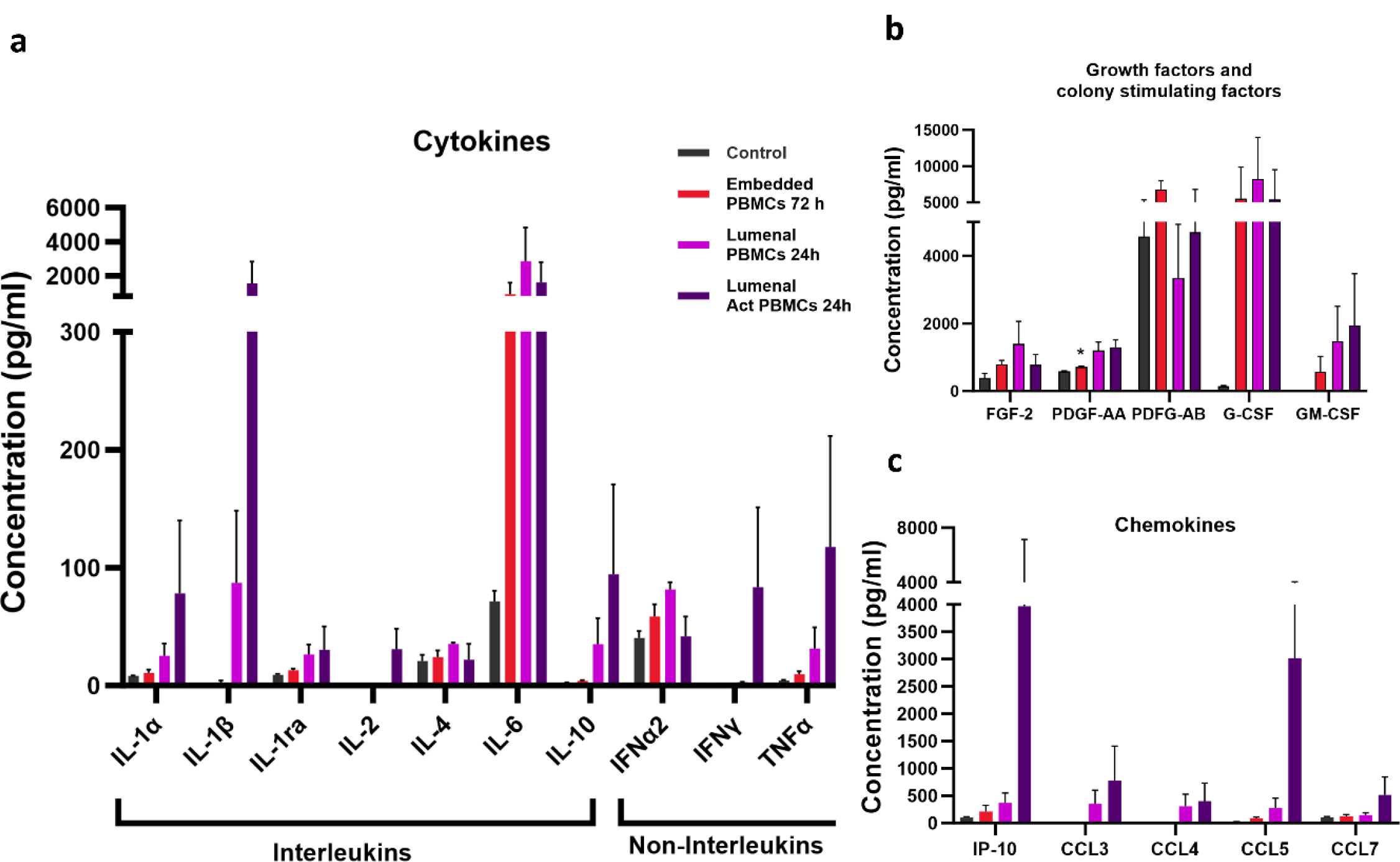
Multiplex analysis of cytokines, colony-stimulating factors, and chemokines from PBMC - HUVEC co-culture in MPS. a - Graph shows average concentration of cytokines in conditioned media collected from HUVEC MPS exposed to PBMCs. b - Graph shows average concentration of growth factors and colony stimulating factors in conditioned media collected from HUVEC MPS exposed to PBMCs. c - Graph shows average concentration of chemokines in conditioned media collected from HUVEC MPS exposed to PBMCs.

## DISCUSSION

### Barrier function

Increased permeability in HUVEC MPS upon treatment with inflammatory mixture is in line with previous studies on lymphatic vessel MPS with IL-6 treatment ^38^ and HUVECs on 2D cultures treated with TNFα ^16^. It is plausible that TNFα is majorly responsible for the poor barrier function ^16^ and likely works via decreased membrane localization of ZO- 1 and VE-cadherin proteins as alluded to in uterine artery endothelial cells ^39^ and in corneal endothelial cells ^40^. Barrier function was improved in HUVEC MPS containing embedded PBMCs, perhaps due to slow and sustained release of relatively low- concentration cytokines that could offer some beneficial effects to the monolayer ^16^. MPS with lumenal activated PBMCs disrupted barrier integrity, possibly explained by the release of higher concentrations of proinflammatory cytokines. This result is consistent with previous reports ^41,42^ A complementary explanation for increased permeability here is that increased chemokine concentrations lead to higher transendothelial migration of PBMCs through the MPS and induce gaps in the monolayer ^43,44^. Overall, our experiments indicate that an inflammatory environment (either from cytokines or activated PBMCs) has negative consequences on endothelial barrier function.

### Secreted factors

Growth factors differed between HUVECs in 2D cultures and 3D MPS (supp tables 1,2) consistent with our previous analysis ^24^. In MPS treated with the inflammatory mixture, multiple other cytokines are secreted supporting the inherent ability of endothelial cells to serve as a source of cytokines ^45^. uPA (urokinase-type plasminogen activator) could potentially explain the observed increase in permeability given that increased microvascular endothelial permeability occurs with uPA treatment ^46^. MPS with lumenal act PBMCs caused increases in cytokines - TNFα and IL-1β compared to MPS with lumenal PBMCs, indicating their involvement in the exacerbated endothelial function between these conditions. Both TNFα and IL-1β have shown to upregulate surface expression of ICAM-1, VCAM-1, E-selectin, P-selectin, CD-40L, and fractalkine (CX3CL1) on endothelial cells ^47–49^, that have implications for interaction and transmigration of lymphocytes ^23^ through the endothelial monolayer.

### Ca^2+^ signaling

Ca^2+^ signaling traditionally follows a biphasic response pattern wherein cells initially respond to ATP stimulation by Ca^2+^ release from intracellular endoplasmic reticulum stores. This initial response is followed by extracellular Ca^2+^ influx ^50,51^ reliant on gap junction communication, mainly through Cx37, Cx40 and Cx43 ^34,52^ in HUVECs. Ca^2+^ network analysis revealed that cells in the MPS were viable considering the 90-100% activity of cells within the first few seconds after ATP addition. Gap-junction communication driven sustained phase Ca^2+^ signaling occurs roughly after 3-4 minutes. In control MPS, 70-75% of cells show coordinated Ca^2+^ activity in this timeframe coinciding with sustained phase Ca^2+^ signaling. This is in agreement with observed synchronization of Ca^2+^ responses in cells from intact Human Umbilical Vein Endothelium ^10^. Interestingly, Ca^2+^ coordination is lost in 2D endothelial cultures. TPA exposure results in decreased gap junction communication ^53,54^. Our result showing decreased co-activity in TPA treated MPS is indeed consistent with previous studies in HUVECs ^34^.

TNFα, VEGF, IL-6 have all individually resulted in reduced Ca^2+^ signaling in HUVECs ^16,17^. As expected, 20-30% of cells showed sustained co-activity in MPS treated with the inflammatory mixture. Interestingly, treatment with the inflammatory mixture for 48h failed to produce any difference in Ca^2+^ co-activity consistent with other studies ^55,56^ likely due to undefined compensatory actions. Our experimental results with CBX are consistent with the ability of CBX to reduce Ca^2+^ signaling in endothelial cells ^57^ and reduce network level Ca^2+^ activity in lymphocytes ^58^. cAMP is a PKA agonist, that promotes gap junction assembly ^37^ and has previously shown to improve Ca^2+^ bursting in uterine artery endothelial cells during a 1h treatment ^59^. As expected, improvement in Ca^2+^ co-activity was observed in HUVEC MPS treated with cAMP 24h + the inflammatory mixture for 1h compared to MPS treated with the inflammatory mixture alone for 1h. This highlights the potential for Ca^2+^ co-activity to be rescued or treated, essential for designing therapies aimed at improving Ca^2+^ response.

The trend towards decreased Ca^2+^ co-activity in MPS interacting with PBMCs hints at the causal role of cytokines secreted from PBMCs. We previously reported that PBMCs cause a decrease in Ca^2+^ signaling in HUVECs, which is rescued upon blocking release of cytokines from PBMCs ^22^. Worsened Ca^2+^ co-activity in MPS with lumenal activated PBMCs is consistent with 2D studies on endothelial cells with activated PBMC interactions ^22^. Overall, our PBMC-HUVEC co-culture experiments are indicative of the ability of PBMCs in long-term interactions with HUVECs to suppress Ca^2+^ signaling in the endothelium.

Our decision to utilize Ca^2+^ co-activity measures is a consequence of recent literature identifying endothelial heterogeneity in Ca^2+^ responses resulting in clusters of cells that show increased responsiveness to Ca^2+^ agonists; in addition to synchronization and network relatedness within these clusters ^60–63^. Synchronized Ca^2+^ responses have in fact recently been measured in HUVECs in a microfluidic device with loss of synchronization upon gap-junction inhibition ^64^. Ca^2+^ network level connectivity has been experimentally validated in endothelial cells isolated from diseased animals; specifically disrupted Ca^2+^ networks were observed in prediabetic obesity ^62^. Importantly, disruption of Ca^2+^ networks was associated with reduced vasodilation, a measure relevant to cardiovascular disorders wherein reduced vasodilation is prevalent. Studies on network activity on endothelial Ca^2+^ signaling including this study have used either intact vessels or cultured cells in microfluidic devices ^62,64^ highlighting the importance of using the 3D MPS system for tissue-level Ca^2+^ analysis. There exists tremendous opportunity to understand cardiovascular diseases and relevant pathophysiology using endothelial cell Ca^2+^ network analyses in MPS models.

## CONCLUSIONS

In summary, we demonstrate that an inflammatory mixture (TNFα, IL-6, VEGF-A - 10ng/ml each) induces endothelial dysfunction in vascular MPS via disrupted network level Ca^2+^ activity with 1h exposure and via increased permeability with 48h exposure. Additionally, lumenal PBMCs in the MPS showed a trend towards dysregulated Ca^2+^ network activity. Activated PBMCs in the MPS led to increased permeability, decline in Ca^2+^ network activity and increased proinflammatory cytokine and chemokine secretion. This MPS model may be leveraged in future for modelling cardiovascular diseases using patient-specific endothelial cells and for testing therapeutics using measures of Ca^2+^ signaling and endothelial permeability.

## AUTHOR CONTRIBUTIONS

AR, MVM, DSB - conceptualized the project and designed experiments. AR, MVM, HEG performed the experiments. AR, MVM, HEG performed data analysis. AR, MVM made figures. DSB, DJB provided funding and resources. AR, MVM and DSB wrote the manuscript, and all authors revised it.

## ACKNOWLEDGEMENTS

School of Medicine and Public Health (SMPH), Department of Obstetrics and Gynecology (Ob-Gyn), Office of the Vice Chancellor for Research and Graduate Education (OVCRGE), Christopher Endemann from Data Science Hub, Social Science Computing Cooperative, L&S Honors Sophomore Summer Apprenticeship at University of Wisconsin-Madison; and Wisconsin Alumni Research Foundation (WARF).

## REFERENCES

1 D. S. Boeldt and I. M. Bird, J. Endocrinol., 2017, 232, R27–R44.

2 G. Gallo, M. Volpe and C. Savoia, Front. Med., 2022, 8, 798958.

3 S. Xu, I. Ilyas, P. J. Little, H. Li, D. Kamato, X. Zheng, S. Luo, Z. Li, P. Liu, J. Han, I. C. Harding, E. E. Ebong, S. J. Cameron, A. G. Stewart and J. Weng, Pharmacol Rev, 2021, 73, 924–967.

4 D. A. Chistiakov, A. N. Orekhov and Y. V. Bobryshev, Front. Physiol.,, DOI:10.3389/fphys.2015.00365.

5 A. Phinikaridou, M. E. Andia, A. Protti, A. Indermuehle, A. Shah, A. Smith, A. Warley and R. M. Botnar, Circulation, 2012, 126, 707–719.

6 Y. Wang, Y. Gu, D. N. Granger, J. M. Roberts and J. S. Alexander, American Journal of Obstetrics and Gynecology, 2002, 186, 214–220.

7 M. S. Taylor, J. Lowery, C.-S. Choi and M. Francis, Front Physiol, 2022, 13, 848681.

8 L. Pogan, L. Garneau, P. Bissonnette, L. Wu and R. Sauvé, J Hypertens, 2001, 19, 721–730.

9 C. Wilson, X. Zhang, C. Buckley, H. R. Heathcote, M. D. Lee and J. G. McCarron, Hypertension, 2019, 74, 1200–1214.

10 J. Krupp, D. S. Boeldt, F.-X. Yi, M. A. Grummer, H. A. Bankowski Anaya, D. M. Shah and I. M. Bird, Am. J. Physiol. Heart Circ. Physiol., 2013, 305, H969–979.

11 A. M. Andrews, T. T. Muzorewa, K. A. Zaccheo, D. G. Buerk, D. Jaron and K. A. Barbee, Cel. Mol. Bioeng., 2017, 10, 30–40.

12 S. Lin, K. A. Fagan, K.-X. Li, P. W. Shaul, D. M. F. Cooper and D. M. Rodman, Journal of Biological Chemistry, 2000, 275, 17979–17985.

13 M. Naya, T. Tsukamoto, K. Morita, C. Katoh, T. Furumoto, S. Fujii, N. Tamaki and H. Tsutsui, Hypertens Res, 2007, 30, 541–548.

14 M. Braile, S. Marcella, L. Cristinziano, M. R. Galdiero, L. Modestino, A. L. Ferrara, G. Varricchi, G. Marone and S. Loffredo, IJMS, 2020, 21, 5294.

15 B. Eržen, M. Šilar and M. Šabovič, BMC Cardiovascular Disorders, 2014, 14, 166.

16 A. K. Mauro, N. Khurshid, D. M. Berdahl, A. C. Ampey, D. Adu, D. M. Shah and D. S. Boeldt, Journal of Endocrinology, 2021, 248, 107–117.

17 A. K. Mauro, D. M. Berdahl, N. Khurshid, L. Clemente, A. C. Ampey, D. M. Shah, M. Bird and D. S. Boeldt, Molecular and Cellular Endocrinology, 2020, 510, 110814.

18 G. Altan-Bonnet and R. Mukherjee, Nat Rev Immunol, 2019, 19, 205–217.

19 K. Thurley, D. Gerecht, E. Friedmann and T. Höfer, PLoS Comput Biol, 2015, 11, e1004206.

20 R. G. Holzheimer and W. Steinmetz, Eur J Med Res, 2000, 5, 347–355.

21 A. E. Norlander, M. S. Madhur and D. G. Harrison, Journal of Experimental Medicine, 2018, 215, 21–33.

22 A. Rengarajan, J. L. Austin, A. K. Stanic, M. S. Patankar and D. S. Boeldt, Reprod. Sci., 2023, 30, 2292–2301.

23 D. Vestweber, Nat Rev Immunol, 2015, 15, 692–704.

24 L. L. Bischel, K. E. Sung, J. A. Jiménez-Torres, B. Mader, P. J. Keely and D. J. Beebe, FASEB j., 2014, 28, 4583–4590.

25 M. Virumbrales-Muñoz, J. M. Ayuso, M. M. Gong, M. Humayun, M. K. Livingston, K. M. Lugo-Cintrón, P. McMinn, Y. R. Álvarez-García and D. J. Beebe, Chem. Soc. Rev., 2020, 49, 6402–6442.

26 K. Gold, A. K. Gaharwar and A. Jain, Biomaterials, 2019, 196, 2–17.

27 V. Paloschi, M. Sabater-Lleal, H. Middelkamp, A. Vivas, S. Johansson, A. van der Meer, M. Tenje and L. Maegdefessel, Cardiovascular Research, 2021, 117, 2742–2754.

28 J. A. Jiménez-Torres, S. L. Peery, K. E. Sung and D. J. Beebe, Adv. Healthcare Mater., 2016, 5, 198–204.

29 D. J. Medina-Leyte, M. Domínguez-Pérez, I. Mercado, M. T. Villarreal-Molina and L. Jacobo-Albavera, Applied Sciences, 2020, 10, 938.

30 V. H. Huxley, F. E. Curry and R. H. Adamson, American Journal of Physiology-Heart and Circulatory Physiology, 1987, 252, H188–H197.

31 T. P. Patel, K. Man, B. L. Firestein and D. F. Meaney, Journal of Neuroscience Methods, 2015, 243, 26–38.

32 J. K. Miller, I. Ayzenshtat, L. Carrillo-Reid and R. Yuste, Proceedings of the National Academy of Sciences, 2014, 111, E4053–E4061.

33 D. R. Stirling, M. J. Swain-Bowden, A. M. Lucas, A. E. Carpenter, B. A. Cimini and A. Goodman, BMC Bioinformatics, 2021, 22, 433.

34 D. S. Boeldt, J. Krupp, F.-X. Yi, N. Khurshid, D. M. Shah and I. M. Bird, American Journal of Physiology-Heart and Circulatory Physiology, 2017, 312, H173–H181.

35 D. S. Boeldt, M. A. Grummer, R. R. Magness and I. M. Bird, Journal of Endocrinology, 2014, 223, 1–11.

36 X. Guan, S. Wilson, K. K. Schlender and R. J. Ruch, Mol. Carcinog., 1996, 16, 157–164.

37 A. F. Paulson, P. D. Lampe, R. A. Meyer, E. TenBroek, M. M. Atkinson, T. F. Walseth and R. G. Johnson, Journal of Cell Science, 2000, 113, 3037–3049.

38 M. M. Gong, K. M. Lugo-Cintron, B. R. White, S. C. Kerr, P. M. Harari and D. J. Beebe, Biomaterials, 2019, 214, 119225.

39 A. C. Ampey, R. L. Dahn, M. A. Grummer and I. M. Bird, Molecular and Cellular Endocrinology, 2021, 534, 111368.

40 M. Shivanna, G. Rajashekhar and S. P. Srinivas, Invest. Ophthalmol. Vis. Sci., 2010, 51, 1575.

41 A. L. B. Seynhaeve, C. E. Vermeulen, A. M. M. Eggermont and T. L. M. ten Hagen, Cell Biochem. Biophys., 2006, 44, 157–169.

42 S. Wakamoto, M. Fujihara, D. Takahashi, K. Niwa, S. Sato, T. Kato, H. Azuma and H. Ikeda, Transfusion, 2011, 51, 993–1001.

43 T. Shigematsu, R. E. Wolf and D. N. Granger, Microcirculation, 2002, 9, 99–109.

44 M. Liu, C.-C. Chien, D. N. Grigoryev, M. T. Gandolfo, R. B. Colvin and H. Rabb, Microvascular Research, 2009, 77, 340–347.

45 G. Krishnaswamy, J. Kelley, L. Yerra, J. K. Smith and D. S. Chi, Journal of Interferon & Cytokine Research, 1999, 19, 91–104.

46 A. M. Makarova, T. V. Lebedeva, T. Nassar, A. A.-R. Higazi, J. Xue, M. E. Carinato, K. Bdeir, D. B. Cines and V. Stepanova, Journal of Biological Chemistry, 2011, 286, 23044–23053.

47 F. Mach, U. Schönbeck, G. K. Sukhova, T. Bourcier, J.-Y. Bonnefoy, J. S. Pober and P. Libby, Proceedings of the National Academy of Sciences, 1997, 94, 1931–1936.

48 G. E. Garcia, Y. Xia, S. Chen, Y. Wang, R. D. Ye, J. K. Harrison, K. B. Bacon, H.-G. Zerwes and L. Feng, J Leukoc Biol, 2000, 67, 577–584.

49 M. Raab, H. Daxecker, S. Markovic, A. Karimi, A. Griesmacher and M. M. Mueller, Clinica Chimica Acta, 2002, 321, 11–16.

50 S. M. Gifford, J. M. Cale, S. Tsoi, R. R. Magness and I. M. Bird, Endocrinology, 2003, 144, 3639–3650.

51 S. P. Srinivas, J. C. Yeh, A. Ong and J. A. Bonanno, Current Eye Research, 1998, 17, 994–1004.

52 H. Van Rijen, M. J. Van Kempen, L. J. Analbers, M. B. Rook, A. C. Van Ginneken, D. Gros and H. J. Jongsma, American Journal of Physiology-Cell Physiology, 1997, 272, C117–C130.

53 S. Sirnes, E. Leithe and E. Rivedal, Biochemical and Biophysical Research Communications, 2008, 373, 597–601.

54 J. L. Solan and P. D. Lampe, Biochimica et Biophysica Acta (BBA) - Biomembranes, 2005, 1711, 154–163.

55 A. C. Ampey, D. S. Boeldt, L. Clemente, M. A. Grummer, F. Yi, R. R. Magness and I. M. Bird, Molecular and Cellular Endocrinology, 2019, 488, 14–24.

56 T. Dalsgaard, S. K. Sonkusare, C. Teuscher, M. E. Poynter and M. T. Nelson, Sci Rep, 2016, 6, 33841.

57 C. Buckley, X. Zhang, C. Wilson and J. G. McCarron, Br J Pharmacol, 2021, 178, 896–912.

58 K. Y. L. Ho, R. J. Khadilkar, R. L. Carr and G. Tanentzapf, Current Biology, 2021, 31, 4697-4712.e6.

59 B. C. Ampey, A. C. Ampey, G. E. Lopez, I. M. Bird and R. R. Magness, Hypertension, 2017, 70, 401–411.

60 M. D. Lee, C. Buckley, X. Zhang, C. Wilson and J. G. McCarron, The FASEB Journal, 2022, 36, fasebj.2022.36.S1.R3354.

61 M. D. Lee, C. Wilson, C. D. Saunter, C. Kennedy, J. M. Girkin and J. G. McCarron, Sci. Signal., 2018, 11, eaar4411.

62 C. Wilson, X. Zhang, M. D. Lee, M. MacDonald, H. R. Heathcote, N. M. N. Alorfi, C. Buckley, S. Dolan and J. G. McCarron, Metabolism, 2020, 111, 154340.

63 C. Wilson, C. D. Saunter, J. M. Girkin and J. G. McCarron, FASEB j., 2016, 30, 2000–2013.

64 A. Zamir, G. Li, K. Chase, R. Moskovitch, B. Sun and A. Zaritsky, Cell Systems, 2022, 13, 711-723.e7.

