## Supplementary material for "Immune Cells and Inflammatory mediators cause endothelial dysfunction in a vascular microphysiological system": supp table

|  | **MPS** | | | | | **2D** |
| --- | --- | --- | --- | --- | --- | --- |
|  | **Control** | **Mix 48h** | **Embedded PBMCs** | **Lumenal PBMCs** | **Lumenal Act PBMCs** | **Control 2D** |
| **FGF-2** | 389.99 | 846.60 | 787.57 | 1398.40 | 781.16 | 0.00 |
| **PDGF-AA** | 582.15 | 568.73 | 716.56 | 1201.82 | 1296.75 | 1156.62 |
| **PDFG-AB** | 4562.75 | 12902.73 | 6745.12 | 3343.60 | 4694.35 | 1232.47 |
| **G-CSF** | 153.43 | 8025.27 | 5514.20 | 8227.04 | 5392.07 | 116.37 |
| **GM-CSF** | 19.76 | 2414.42 | 573.25 | 1473.37 | 1941.74 | 2.81 |
| **CXCL1** | 11129.86 | 11188.95 | 10898.40 | 6638.38 | 10639.63 | 6602.65 |
| **CXCL10** | 103.23 | 12376.98 | 211.06 | 369.54 | 3970.14 | 60.62 |
| **CX3CL1** | 43.60 | 1389.57 | 65.39 | 73.94 | 185.67 | 35.65 |
| **CCL2** | 9342.40 | 8794.89 | 9972.42 | 13872.36 | 9508.84 | 5076.56 |
| **CCL3** | 1.37 | 3.32 | 6.94 | 353.36 | 775.90 | 0.36 |
| **CCL4** | 1.22 | 2.23 | 3.47 | 310.71 | 404.90 | 0.80 |
| **CCL5** | 20.80 | 3254.19 | 89.73 | 273.67 | 3019.76 | 10.30 |
| **CCL7** | 104.86 | 695.57 | 122.95 | 146.91 | 513.71 | 35.70 |
| **CCL22** | 8.18 | 12.72 | 13.31 | 20.44 | 50.61 | 6.56 |
| **TNFα** | 4.29 | 8853.45 | 9.57 | 31.47 | 117.68 | 1.19 |
| **IFNα2** | 40.48 | 50.35 | 58.45 | 81.42 | 41.87 | 36.36 |
| **IFNγ** | 1.39 | 1.69 | 1.83 | 3.05 | 83.38 | 1.19 |
| **IL-1RA** | 9.00 | 12.82 | 12.99 | 26.43 | 30.42 | 7.20 |
| **IL-1a** | 8.23 | 14.80 | 10.82 | 25.36 | 78.02 | 4.75 |
| **IL-1β** | 0.25 | 0.95 | 2.55 | 87.15 | 1563.65 | 0.19 |
| **IL-2** | 0.58 | 1.16 | 1.03 | 0.92 | 30.79 | 0.37 |
| **IL-4** | 20.91 | 34.29 | 24.20 | 35.43 | 21.97 | 13.48 |
| **IL-6** | 71.32 | 4276.20 | 925.45 | 2857.18 | 1628.46 | 143.60 |
| **IL-10** | 2.42 | 3.10 | 3.92 | 34.99 | 94.39 | 1.96 |

Supplementary Table 1 – Multiplexed analysis of secretion media from vascular MPS and 2D cultures.

Concentrations of different factors are listed in ng/ml.

|  | **MPS** | | | | |  |
| --- | --- | --- | --- | --- | --- | --- |
| **Fold difference** | **Mix 48h vs Control** | **Embedded PBMCs vs Control** | **Lumenal PBMCs vs Control** | **Lumenal Act PBMCs vs Control** | **Lumenal Act PBMCs vs Lumenal PBMCs** | **Control 2D vs 3D** |
| **FGF-2** | 2.17 | 2.02 | 3.59 | 2.00 | 0.56 | 0.00 |
| **PDGF-AA** | 0.98 | 1.23 | 2.06 | 2.23 | 1.08 | 1.99 |
| **PDFG-AB** | 2.83 | 1.48 | 0.73 | 1.03 | 1.40 | 0.27 |
| **G-CSF** | 52.31 | 35.94 | 53.62 | 35.14 | 0.66 | 0.76 |
| **GM-CSF** | 122.18 | 29.01 | 74.56 | 98.26 | 1.32 | 0.14 |
| **CXCL1** | 1.01 | 0.98 | 0.60 | 0.96 | 1.60 | 0.59 |
| **CXCL10** | 119.90 | 2.04 | 3.58 | 38.46 | 10.74 | 0.59 |
| **CX3CL1** | 31.87 | 1.50 | 1.70 | 4.26 | 2.51 | 0.82 |
| **CCL2** | 0.94 | 1.07 | 1.48 | 1.02 | 0.69 | 0.54 |
| **CCL3** | 2.42 | 5.07 | 258.11 | 566.75 | 2.20 | 0.26 |
| **CCL4** | 1.83 | 2.84 | 254.61 | 331.80 | 1.30 | 0.65 |
| **CCL5** | 156.44 | 4.31 | 13.16 | 145.17 | 11.03 | 0.50 |
| **CCL7** | 6.63 | 1.17 | 1.40 | 4.90 | 3.50 | 0.34 |
| **CCL22** | 1.56 | 1.63 | 2.50 | 6.19 | 2.48 | 0.80 |
| **TNFα** | 2066.03 | 2.23 | 7.34 | 27.46 | 3.74 | 0.28 |
| **IFNα2** | 1.24 | 1.44 | 2.01 | 1.03 | 0.51 | 0.90 |
| **IFNγ** | 1.22 | 1.32 | 2.20 | 60.11 | 27.32 | 0.86 |
| **IL-1RA** | 1.42 | 1.44 | 2.94 | 3.38 | 1.15 | 0.80 |
| **IL-1a** | 1.80 | 1.32 | 3.08 | 9.48 | 3.08 | 0.58 |
| **IL-1β** | 3.80 | 10.15 | 347.13 | 6228.13 | 17.94 | 0.76 |
| **IL-2** | 2.00 | 1.76 | 1.58 | 52.82 | 33.40 | 0.64 |
| **IL-4** | 1.64 | 1.16 | 1.69 | 1.05 | 0.62 | 0.64 |
| **IL-6** | 59.95 | 12.98 | 40.06 | 22.83 | 0.57 | 2.01 |
| **IL-10** | 1.28 | 1.62 | 14.47 | 39.03 | 2.70 | 0.81 |

Supplementary Table 2 – Fold differences in concentration of secretion media from vascular MPS and 2D cultures.
